# Genome-wide identification and analyses of tomato (*Solanum lycopersicum* L.) high-affinity nitrate transporter 2 (*NRT2*) family genes and their responses to drought and salinity

**DOI:** 10.1101/2021.12.08.471831

**Authors:** M. Aydın Akbudak, Ertugrul Filiz, Durmuş Çetin

## Abstract

High-affinity nitrate transporter 2 (NRT2) proteins have vital roles in nitrate (NO_3_^-^) uptake and translocation in plants. The gene families coding NRT2 proteins have been identified and functionally characterized in many plant species. However, no systematic identification of NRT2 family members have been reported in tomato (*Solanum lycopersicum*). There is also little known about their expression profiles under environmental stresses. Accordingly, the present study aimed to identify NRT2 gene family in the tomato genome; then, investigate them in detail through bioinformatics, physiological and expression analyses. As a result, four novel *NRT2* genes were identified in the tomato genome, all of which contain the same domain belonging to the Major Facilitator Superfamily (PF07690). The co-expression network of *SlNRT* genes revealed that they were co-expressed with several other genes in many different molecular pathways including transport, photosynthesis, fatty acid metabolism and amino acid catabolism. Programming many crucial physiological and metabolic pathways, various numbers of phosphorylation sites were predicted in the NRT2 proteins.

## Introduction

Nitrogen (N) is an essential macronutrient, constituting important cellular molecules such as amino acids, nucleic acids, chlorophyll (Hawkesford et al., 2012). Approximately 170 million tons of nitrogenous fertilizer, mainly in the nitrate and ammonium forms, is used for agricultural production in the world every year (FAO, 2019). However, 50 – 70% of this costly input were not taken up by plants and rests in the soil (Hodge et al., 2000; Gramma et al., 2020). The excess nitrogen, then, infiltrates into the underground water, causing various environmental and health issues. Therefore, optimizing the amounts of the nitrogenous fertilizers to be used and improving the nitrogen use efficiency of the plants are two of the main goals of breeders and scientists working in the plant nutrition area.

Nitrate (NO_3_^-^) is the most abundant form of inorganic nitrogen in the soil, and its uptake and translocation in plants play crucial roles in nitrogen use efficiency (Jin et al., 2015). It is actively taken up through roots and leaves, and transported in the plant by nitrate transporters, each of which has different features (Bai et al., 2013; Guan, 2017). NITRATE TRANSPORTER 1 (NRT1) / PEPTIDE TRANSPORTER (PTR) family (NPF), NRT2, CHLORIDE CHANNEL (CLC) and slowly activating anion channel protein family are the four protein families which function in nitrate transport (Fan et al., 2017; Wang et al., 2018).

Nitrate assimilation is energy-costly, because it requires ATP, reducing equivalents and carbon (C) skeletons usage in high amounts (Nunes-Nesi et al., 2010). Hence, it is subjected to be regulated according to the availability of nitrogen sources in the environment and plant needs changing in compliance with the developmental stages. Plant roots have two different nitrate transport systems, LATS and HATS. The LATS (Low Affinity Transport System) is activated by high nitrate concentrations (> 1 mM) while the HATS (High Affinity Transport System) gets activated by low (<1 mM) nitrate concentrations in the soil (Dechorgnat et al., 2012). Besides plant nutrient requirements, environmental factors such as drought and salinity regulate the activation and deactivation of these systems (Yao et al., 2008). To understand the mechanism controlling nitrate uptake and distribution in the plants better, identification of the genes and the proteins responsible for nitrate transport is crucial.

Research showed that NRT1 proteins function as the main component of the LATS activated in high nitrate concentrations. On the other hand, NRT2 family proteins function in the HATS system under low nitrate concentrations (Tsay et al., 2007). Still, some NRT1 proteins have dual function in both HATS and LATS systems. Despite sharing the same 3-D structure, there is no primary sequence homology between NRT1 and NRT2 proteins (Orsel et al., 2002). They have at least 2 members per plant species. Many of the NRT2 proteins were found in the root tissue; however, the information is limited if they transport nitrate through the plant and between the cell compartments.

The presence of *NRT* genes in high numbers and their wide-distribution in the genome reveal their importance for the plant life cycle. 178 *NRT1*, 20 *NRT2* and 6 *NRT3* gene have been identified in *Saccharum spontaneum* genome, as distributed to its all eight chromosomes (Wang et al. 2020). 120 *NRT1* and 5 *NRT2* genes have been identified in wild soybean (*Glycine soja*) (You et al., 2020). Their gene structure and protein motifs revealed that GsNRT family members were conserved in both genomic and peptide sequences. They function not only in plant growth and development but also in adaptation to environmental stresses.

The first identified eukaryotic *NRT2* gene, *crnA* was isolated from a filamentous fungus,*Aspergillus nidulans* approximately 30 years ago (Johnstone et al., 1990; Unkless et al., 1991). Using their sequence homologies with *crn*A, some barley (Trueman et al., 1996), *Nicotiana plumbaginifolia* (Quesada et al., 1997), soybean (Amarashinghe et al., 1998) and tomato (Ono et al., 2000) *NRT2* genes have been identified and functionally characterized. In the present study, genome-wide identifications and comprehensive bioinformatics analyses of *NRT2* genes were performed in tomato (*Solanum lycopersicum, Sl*). This is also the very first study presenting *SlNRT2* genes expression profiles in response to drought stress.

## Materials and method

### Identification of *NRT2* genes in tomato

The protein sequences of *Arabidopsis* NRT2 (AT1G08090, AT1G08100, AT5G60780, AT5G60770, AT1G12940, AT3G45060 and AT5G14570) were obtained from UniProt database (https://www.uniprot.org/) and these sequences were used as references sequences for blastp analyses in tomato (*Solanum lycopersicum*, Sl) genome in Phytozome v12.1 database (https://phytozome.jgi.doe.gov/pz/portal.html#!info?alias=Org_Slycopersicum). Downloaded from Pfam database (http://pfam.xfam.org/), the Hidden Markov Model (HMM) profiles of the MSF_1 (PF07690) domain were used to confirm the core domain. Finally, all non-redundant protein sequences were accepted as tomato NRT2 proteins. ProtParam server (https://web.expasy.org/protparam/) was used to compute various physical and chemical parameters of SlNRT2 protein sequences, including their molecular weights, isoelectric points (*pI*) and protein lengths (Gasteiger et al. 2005). The exon/intron structures were constructed using GSDS (http://gsds.cbi.pku.edu.cn/) (Hu et al. 2015). CELLO v.2.5 server (http://cello.life.nctu.edu.tw/) was used to predict the subcellular localization of NRT2 proteins in tomato (Yu et al. 2006). Prediction of transmembrane helices in SlNRT2 proteins was evaluated using TMHMM 2.0 online server (http://www.cbs.dtu.dk/services/TMHMM/) (Krogh et al. 2001). KEGG annotation analyses were performed using KEGG online server such as KEGG pathway, KEGG module and KEGG orthology (https://www.genome.jp/kegg/).

### Motif and phylogenetic analyses

Firstly, the multiple sequence alignment and sequence identity matrix of the proteins were constructed by BioEdit v7.2.5 software (Hall, 1999). The conserved motifs were investigated using Multiple Expectation maximization for Motif Elicitation (MEME v5.3.3) algorithm in MEME suite of analysis tools (https://meme-suite.org/meme/tools/meme) (Bailey et al., 2015). The parameters for the search were adopted as: max motif number to find: 10 and min–max motif width to find: 6–50. The molecular phylogenetic analysis of SlNRT2 proteins was evaluated using MEGAX v10.1.8 (Kumar et al., 2018). The evolutionary history was inferred using the maximum likelihood method (ML), based on the JTT matrix-based model. The bootstrap consensus tree was constructed from 1000 replicates by applying neighbor-join and BioNJ algorithms.

### Digital *SlNRT2* gene expression, co-expression, promotor and synteny analyses

To conduct the digital gene expression and co-expression network analyses, the expression data of *S. lycopersicum* tissues and organs (seed, root, meristem, leaf, flower and fruit) were obtained from the TomExpress platform (http://tomexpress.toulouse.inra.fr) (Zouine et al. 2017). The promoter sequences of *SlNRT2* genes were analyzed using the 1500 bp region upstream from the start codon (ATG) for each gene. They were obtained from the Phytozome Database (https://phytozome.jgi.doe.gov/pz/portal.html#!info?alias=Org_Slycopersicum), and submitted to plantCARE (http://bioinformatics.psb.ugent.be/webtools/plantcare/html/) server (Lescot et al. 2002). The synteny analyses of tomato, *Arabidopsis* and rice *NRT2* genes were investigated using the Circos software package (http://circos.ca/).

### Prediction of 3D structure and phosphorylation sites

The secondary structure analyses of SlNRT2 proteins were conducted using SOPMA server, including helix, sheet, turn and coil (https://npsa-prabi.ibcp.fr/cgi-bin/npsa_automat.pl?page=/NPSA/npsa_sopma.html) (Geourjon and Deleage 1995). Putative tertiary structures were generated using the Protein Homology/analogy Recognition Engine V 2.0 (Phyre2) server (http://www.sbg.bio.ic.ac.uk/~phyre2/html/page.cgi?id=index) (Kelley et al., 2015). The molecular cavities and their geometric properties were predicted using BetaCavityWeb online server (http://voronoi.hanyang.ac.kr/betacavityweb/) (Kim et al. 2015). The serine, threonine or tyrosine phosphorylation sites were identified using the NetPhos 3.1 server (http://www.cbs.dtu.dk/services/NetPhos/) (Blom et al. 1999). N-Glycosylation and SUMOylation sites were predicted using NetNGlyc 1.0 (http://www.cbs.dtu.dk/services/NetNGlyc/) and GPS-SUMO (http://sumosp.biocuckoo.org/) online servers, respectively (Gupta and Brunak, 2001; Zhao et al. 2014). In addition, the lysine acetylation and palmitoylation sites were predicted using GPS-PAIL (http://pail.biocuckoo.org/) and CSS-Palm (http://csspalm.biocuckoo.org/) servers, respectively (Ren et al. 2008; Deng et al. 2016).

### Plant materials and stress treatments

*S. lycopersicum* (Adel, F1) plants were grown in peat: perlite (3:1) mixture at 25 °C and 50% humidity under 16h photoperiod and 140 μmol m^−2^ s^−1^ PPFD. They were watered with ½ Hoagland solution water every other day. While control plants were continued to be watered, watering was entirely stopped with the drought-stressed plants after 40 days of plant growth. Seven-days of drought treatment resulted in wilting, and plant leaves were harvested for RNA and protein isolation.

To induce salinity stress, the plants were watered with tap water containing supplemental sodium chloride (NaCl). Salt concentration was increased gradually (50mM, 100mM and 200mM) every 48h intervals to avoid osmotic shock (Swami et al., 2011) Upon 12h treatment with 200mM NaCl, leaves were harvested for RNA and protein isolation and physiological assays.

### MDA (malondialdehyde), proline and H_2_O_2_ assays

The MDA content of plants was assayed using a method modified from Ohkawa et al. (1979). Grounded in liquid nitrogen, 0.2 g of leaf or root tissue was suspended in 2 ml 5% Trichloroacetic acid (TCA). The homogenates were transferred into clean 2 ml microfuge tubes and centrifuged at 12000 rpm in room temperature. Equal amounts of lysate and freshly prepared 0.5 % thiobarbituric asit (TBA) in 20% TCA were mixed, and then incubated at 96°C for 25 min. The tubes were chilled on ice till reaching the room temperature, and centrifuged at 10000 rpm for 5 min. The absorbances of the supernatants were measured at 532 nm and 600 nm wavelengths to clear off non-specific reflections due to turbidity. A freshly prepared 0.5% TBA in 20%TCA solution was used as blank. The MDA content of the samples was assayed using an absorbance coefficient of 155 mM^−1^ cm^−1^.

To assay the proline content of the tissues, 0.5 g of leaf or root tissue was grounded in liquid nitrogen, and suspended in 10 ml of 3% (v/v) sulfosalicylic acid. The suspensions were filtered out through filter papers (Whatman, USA). 2 ml of each filtrate were mixed with 2 ml acid ninhydrin and 2 ml glacial acetic acid, and kept in 100 °C for one hour. The reaction was terminated by placing the mixes onto ice, and 4 ml of toluene was added into each. The chromophore phases consisted were transferred into a quartz plate and the absorbances were measured at 520 nm using a microplate spectrophotometer (Thermo Fischer Scientific, USA).

The hydrogen peroxide contents was assayed using the method developed by Sergiev et al. (Sergiev et al. 1997). After grounded in liquid nitrogen, 0.5 g of a plant sample was homogenized in 5 ml 0.1% trichloroacetic acid (TCA). Following the centrifugation at 12000 g for 15 minutes, 0.5 ml supernatants were transferred into clean 1.5 ml microfuge tubes, and 0.5 ml 10 mM potassium phosphate buffer (pH 7.0) and 1 ml 1 M potassium iodide (KI) were added into them. The absorbances of the supernatants were measured at 390 nm. The H_2_O_2_ contents of the samples were calculated using a standard curve, which was prepared according to known H_2_O_2_ concentrations.

### RNA isolation and gene expression analysis

RNA from leaf and root tissues were isolated using RNA Plant Mini Kit (Qiagen, Cat No: 74904) according to the manufacturer’s instructions. RNA samples were treated with RQ1 RNase-Free Dnase (Promega, USA). The intactness of RNA and presence of DNA contamination in the samples were checked by gel electrophoresis. The RNA amounts were determined using Qubit (Invitrogen, USA). RT-qPCR was carried out in Light Cycler 96 System (Roche). The expressions of the genes were quantified in 10 ng RNA samples using Luna Universal One-Step RT-qPCR Kit (NEB, USA). The forward and reverse primers (Table 1) of the genes were designed for RT-qPCR analysis. As an endogenous control, actin isoform B (*Actin*) gene was used as reference (Goupil et al., 2009)

**Table 1.**
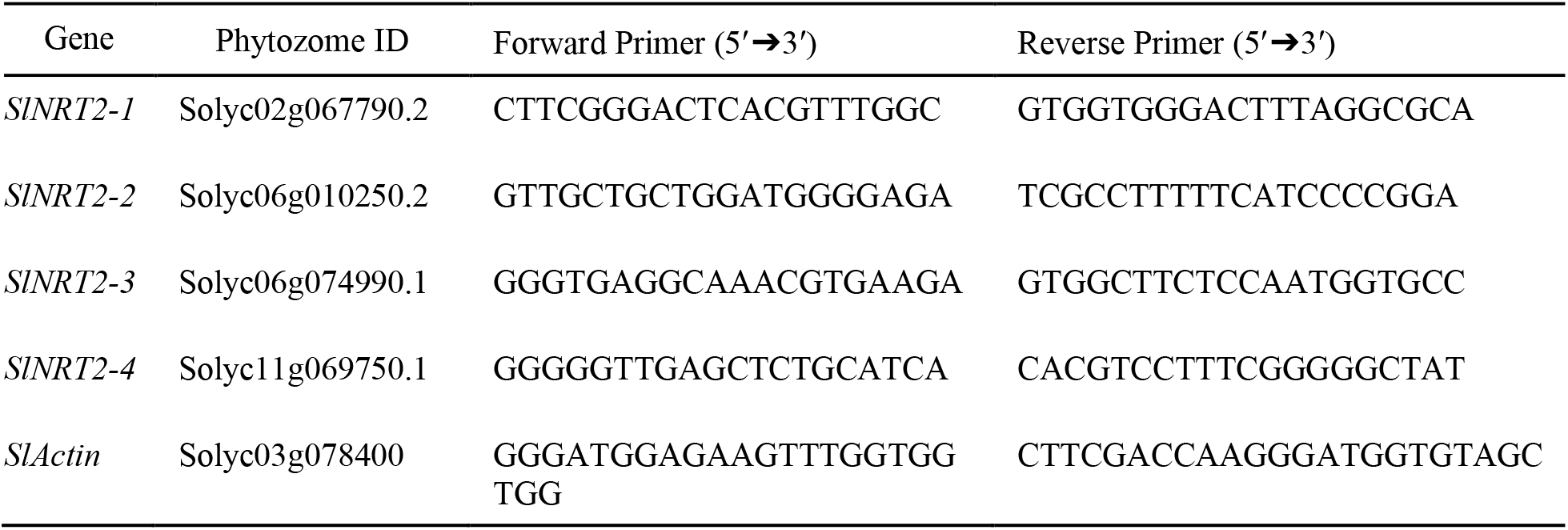
Primers used for RT-qPCR analysis of the *SlNRT2* genes.

## Results and discussion

### Identification of *NRT2* genes in the tomato genome

Firstly, four *SlNRT2* genes were identified using bioinformatics analyses and compared the *Arabidopsis* and rice NRT2 sequences in Table 1. Recent studies also revealed the presence of *NRT2* genes in several other plant species, including 20 *NRT2* genes in wild sugarcane (Wang et al., 2020), 17 in canola (Tong et al., 2020), seven in *Arabidopsis* (Orsel et al., 2002), six in rice (Cai et al., 2008), six in poplar (Bai, 2013), six in cassava (You et al., 2021), five in wild soybean (You et al., 2020), five in apple (Tahir et al., 2021) and four in barley (Guo et al., 2020) have been identified. All NTR2 proteins contain PF07690 (Major Facilitator Superfamily) domain structure and protein length ranged from 460 to 539 amino acid residues. While molecular weight values were found between 50.00 and 58.64 kDa, subcellular localizations were predicted as plasma membrane for all NRT2 proteins. In addition, all NRT2 proteins showed the alkaline characteristics based on *pI* values, except from AT5G14570 (NRT2.7). Similar results were identified in *G. soja* (You et al. 2020). The number of transmembrane helix was ranged from nine to 11, proving transport activity of *NRT2* genes. Wang et al. (2019) reported that *NRT2* genes only have one or two exons, except for SsNRT2.19 in *Saccharum spontaneum*. In *G. soja*, five *GsNRT2* gene found as two or three exons (You et al. 2020). In this study, the exon number of *NRT2* genes showed the high variation level between one and four, suggesting that these results may prove the functional diversities of *NRT2* genes in tomato.

**Table 1.** The general information and sequence details of tomato, *Arabidopsis*, and rice NRT2 genes/proteins.

| Phytozome ID | Species | Exon no | Protein length (aa) | Domain family | Mol. wt.(kDa) | *pI* | SL | TMH |
| --- | --- | --- | --- | --- | --- | --- | --- | --- |
| Solyc02g067790.2 (NRT2.1) | *Sl* | 2 | 460 | PF07690 | 50.00 | 9.04 | PM | 9 |
| Solyc06g010250.2 (NRT2.2) | *Sl* | 4 | 498 | PF07690 | 54.47 | 9.10 | PM | 10 |
| Solyc06g074990.1 (NRT2.3) | *Sl* | 5 | 526 | PF07690 | 57.91 | 9.78 | PM | 10 |
| Solyc11g069750.1 (NRT2.4) | *Sl* | 3 | 525 | PF07690 | 57.49 | 9.05 | PM | 11 |
| AT1G08090 (NRT2.1) | *At* | 3 | 530 | PF07690 | 57.71 | 8.85 | PM | 11 |
| AT1G08100 (NRT2.2) | *At* | 3 | 522 | PF07690 | 56.61 | 8.72 | PM | 11 |
| AT5G60780 (NRT2.3) | *At* | 3 | 539 | PF07690 | 58.23 | 8.96 | PM | 11 |
| AT5G60770 (NRT2.4) | *At* | 4 | 527 | PF07690 | 57.77 | 9.04 | PM | 11 |
| AT1G12940 (NRT2.5) | *At* | 3 | 502 | PF07690 | 54.26 | 8.97 | PM | 11 |
| AT3G45060 (NRT2.6) | *At* | 2 | 542 | PF07690 | 58.64 | 8.84 | PM | 11 |
| AT5G14570 (NRT2.7) | *At* | 2 | 493 | PF07690 | 52.68 | 5.40 | PM | 10 |
| LOC_Os01g50820.1 | *Os* | 1 | 516 | PF07690 | 55.41 | 8.99 | PM | 10 |
| LOC_Os02g02190.1 | *Os* | 1 | 533 | PF07690 | 57.23 | 8.43 | PM | 11 |
| LOC_Os01g36720.1 | *Os* | 2 | 485 | PF07690 | 50.20 | 8.37 | PM | 10 |

PF07690: Major Facilitator Superfamily, PM: plasma membrane (CELLO server), TMH: transmembrane helix

To get more insights about NRT2 proteins, sequence identity matrix was constructed for tomato, rice, and *Arabidopsis* (data not shown). Based on identity data, it was determined that the SlNRT2.1 protein was most similar to AtNRT2.7 with 59.9%. Remarkably, SlNRT2.2, 2.3, and 2.4 proteins showed the greatest similarity with AtNRT2.4 protein with 67.4%, 65.5%, and 69.3%, respectively. KEGG annotation analyses of *SlNRT2* genes were performed using KEGG online server, including KEGG pathway, KEGG module, and KEGG orthology. According to KEGG database’s results, nitrogen metabolism (sly00910) for pathway, nitrate assimilation (sly_M00615) for KEGG Module, and MFS (Major Facilitator Superfamily) transporter, NNP family, and nitrate/nitrite transporter for the KO (KEGG Orthology) definition were identified, proving roles of *SlNRT2* genes in nitrogen assimilation.

### Conserved motif and phylogenetic analyses

10 conserved motifs were identified using MEME server and motif 1, 2, 3, 4, 6 and 10 were present in all NRT2 protein sequences (Fig. 1). Moreover, motif 7 (blue) and motif 8 (purple) were not present in all NRT2s. Remarkably, no relationship of any motif structure with domain sequences has been determined. In general terms, it can be said that there are conserved motif structures for NRT2 proteins.

**Fig. 1.**
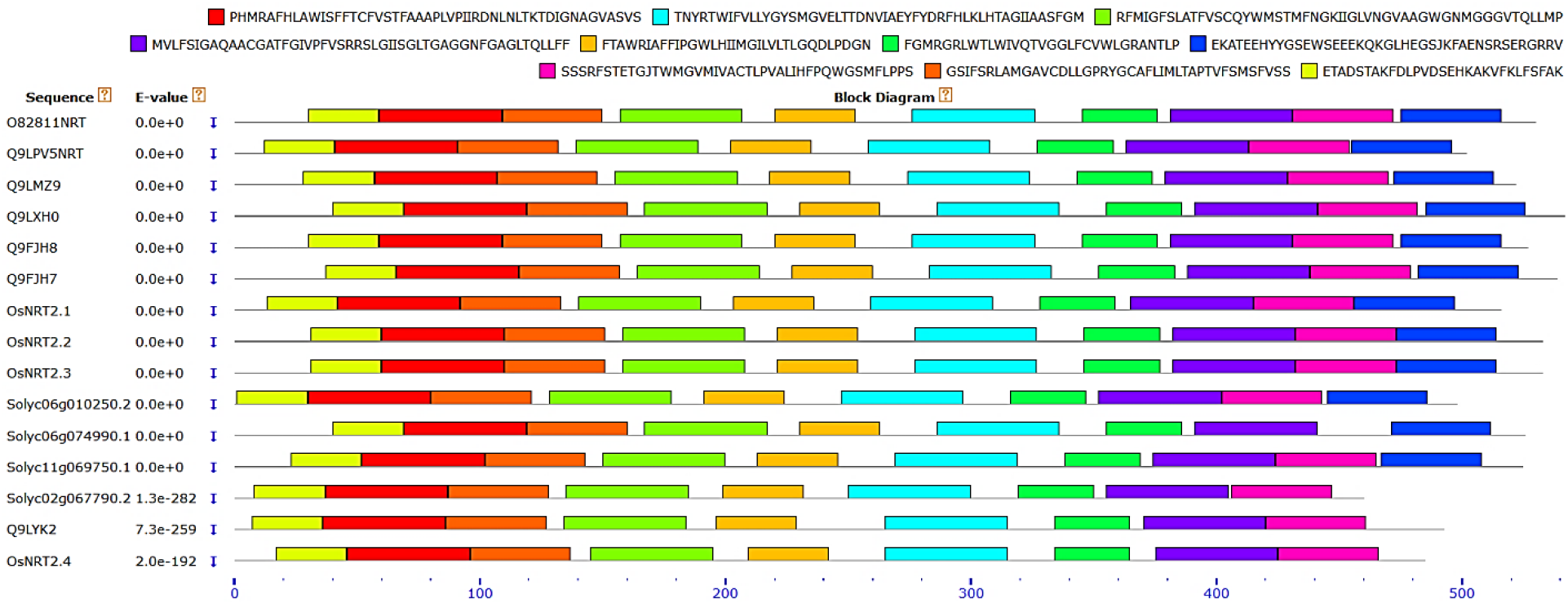
The block diagram of 10 conserved motifs of tomato, *Arabidopsis*, and rice NRT2 sequences using MEME server. In addition, each colors represent the different motif structure and position.

To get more insights about evolutionary history, phylogenetic analyses was performed from tomato, *Arabidopsis* and rice NRT2 protein sequences using MEGAX with NJ method (Fig. 2). Two major groups were identified such as group A with subgroup A1 and A2 and group B. Remarkably, monocot and dicots species were clustered together in subgroup A1, whereas subgroup A2 and group B consist of only dicot species. Similar results were found in phylogenetic analyses of barley NRT2 proteins (Guo et al. 2020). The co-clustering of dicots and monocot may be related to the high level of interspecies conservation of *NRT2* genes in plants.

**Fig. 2.**
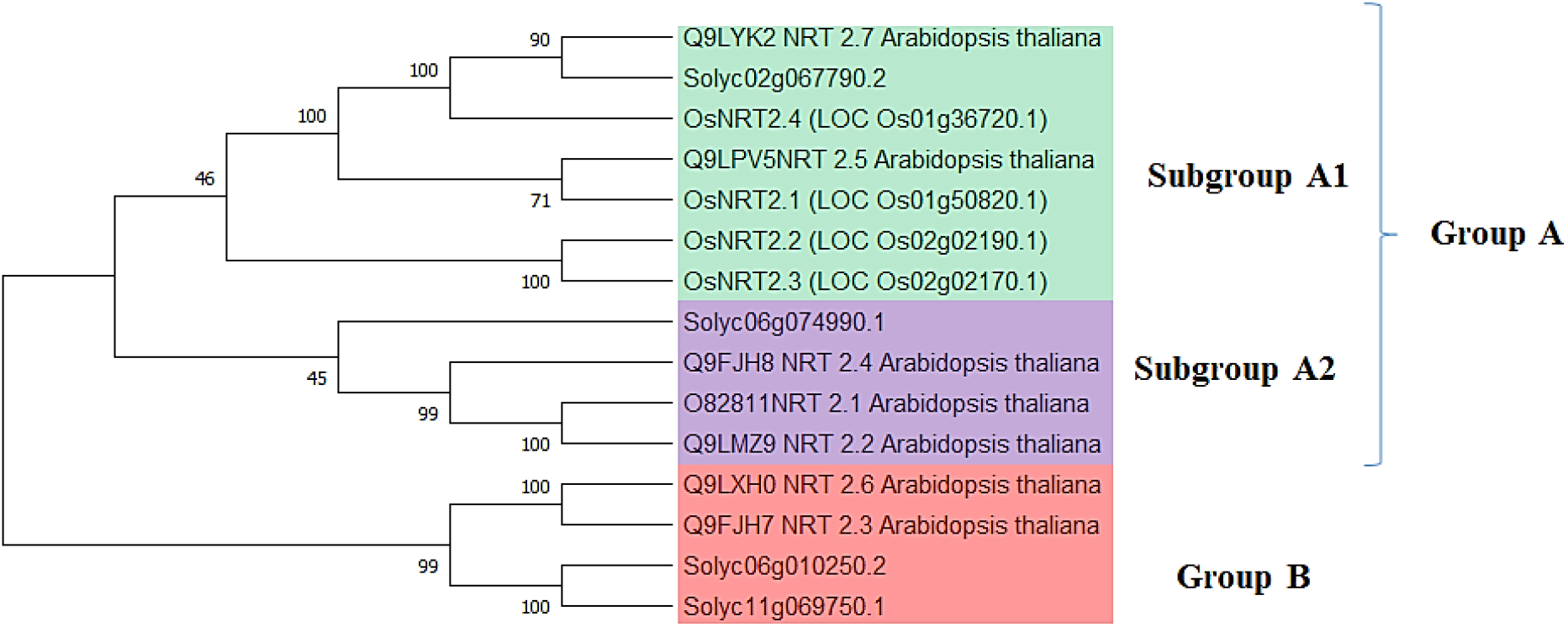
Phylogenetic tree of the NRT2s of tomato, Arabidopsis and rice using NJ tree with 1000 bootstrap values. In addition, each colors represent the different group or subgroup in phylogenetic tree.

### Analysis of the *cis*-acting elements

To obtain the *cis*-acting regulatory elements (CREs) of the *SlNRT2* gene family, we analyzed promotor sequences using Plant-CARE online server (Fig. 3). A total of 11 types CRES were identified and these regulatory elements mainly include hormone response, including abscisic acid, salicylic acid, gibberellin, methyl jasmonate and auxin. In apple, all NRT2 and NRT3 subfamilies contain jasmonic acid and abscisic acid motifs except MdNRT2.4 and MdNRT2.5 respectively (Tahir et al. 2021). Tong et al. (2020) stated that CREs of rapeseed *NRT2* genes can be grouped four categories such as abiotic stress responsive elements, development regulative elements, *MYB* binding sites and hormone-related elements. Also, it was reported that phytohormones play roles in N regulation and signaling, including auxin, cytokinin and both abscisic acid and brassinosteroid (Kiba et al. 2011). Broad spectrum CREs owned by *NRT2* genes may support this gene family to play an active role in cellular metabolic pathways, particularly hormone signaling pathways.

**Fig. 3.**
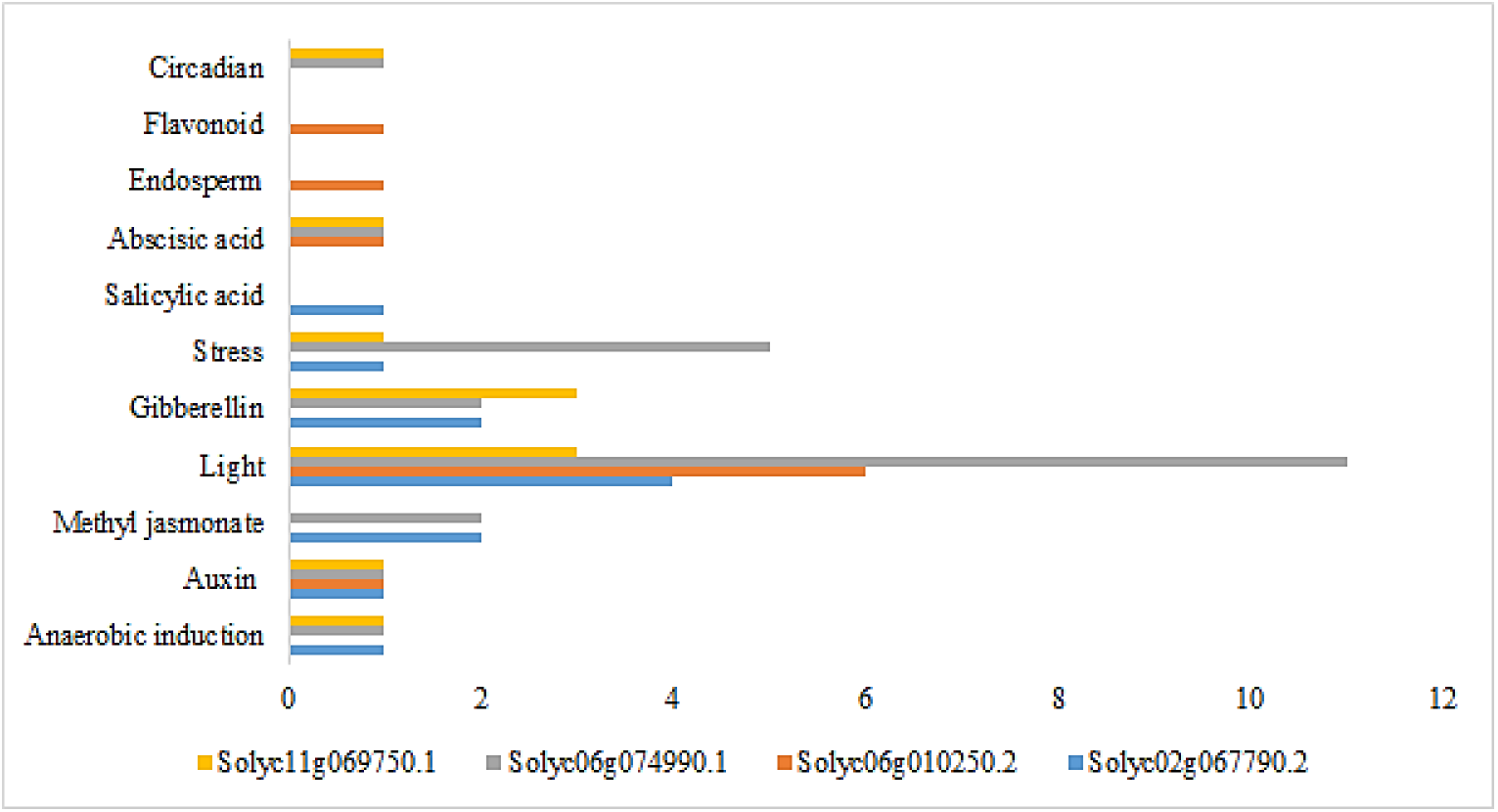
Predicted *cis*-elements in the promoters of the *SlNRT2* genes. The 1500 bp sequences of four *SlNRT2* genes were analyzed by the Plant-CARE online server

### Digital gene expression profile and synteny analysis of *SlNRT2s*

Digital gene expression analyses of *SlNRT2* genes were obtained in different organs and tissue types from TomExpress platform (Fig. 4). In general, *SlNRT2.1* and *SlNRT2.2* genes showed the higher expression levels that other *SlNRT2* genes, whereas *SlNRT2.4* gene indicated the lowest expression level in many tissues and organs. Remarkably, while *SlNRT2.3* gene showed the highest expression level in root, *SlNRT2.1* gene was the highest expression level in anthesis stage. In barley, *HvNRT2* genes generally showed low expression level in eight selected tissues from digital RNA-seq data. However, roots from seedling, 4 days embryos and roots (28 DAP) indicated the higher expression level than other tissues (Guo et al. 2020). In rapeseed, *BnNRT2* genes showed higher expression level in root, flower, leaf and pericarp tissues based on digital RNA-seq analysis (Tong et al. 2020). In poplar, *PtNRT2* genes showed higher expression level in leaves, wood and root tissues based on digital RNA-seq analysis (Bai et al. 2013). In general, it can be said that *SlNRT2* genes play an active role in different tissues, organs and developmental stages.

**Fig. 4.**
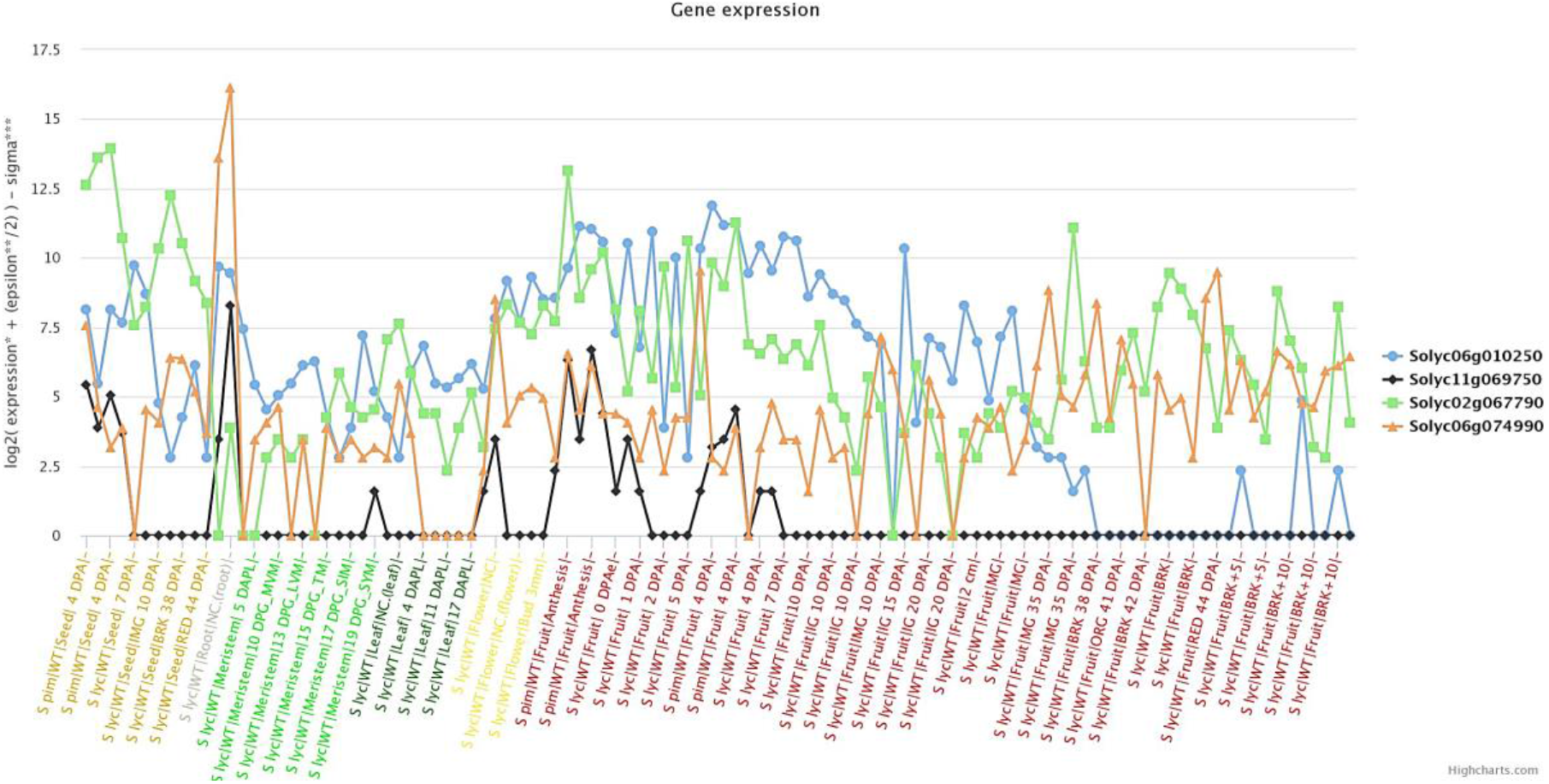
Expression profiles of four *SlNRT2* genes at different developmental stages and tissue types using TomExpress platform

Comparative genomics can contribute to understanding of evolutionary relationship of different species. In this study, synteny analysis was constructed using tomato, *Arabidopsis* and rice *NRT2* genes by Circos tool (Fig. 5). It was found that tomato and *Arabidopsis* species were mostly connected by red lines with more than 90% identity. Also, when tomato and rice were examined, identity ratios of syntenic genes were found to be lower. As a result, synteny-based analysis indicated that more similar syntheny blocks were identified between tomato and *Arabidopsis*, suggesting they have common ancestor before functional divergence in their evolutionary history.

**Fig. 5.**
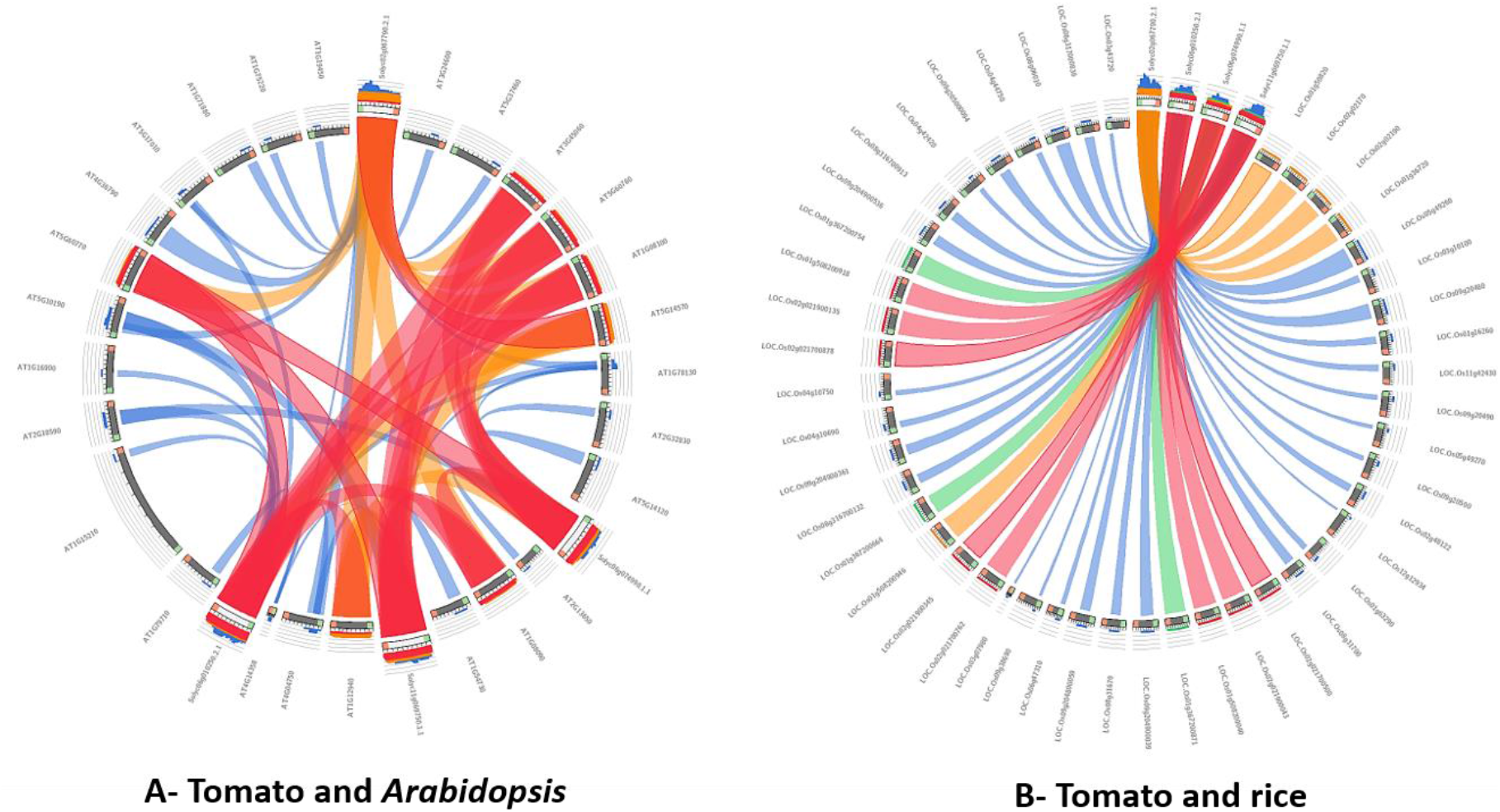
The synteny relationship of *NRT2* gene family in tomato, *A. thialana*, and rice by using the Circos tool. The blue, green, orange and red colors show ≤ 50%, ≤ 70%, ≤ 90% and >90% identity, respectively.

### Co-expression network analyses of *SlNRT2* genes

The co-expression network analyses were constructed for each *SlNRT2* genes using TomExpress platform under abiotic stress conditions (Fig. 6). In this context, five top genes that showed the highest correlation score with *SlNRT2* genes were identified and annotated. For *SlNRT2.1* gene, Solyc02g082200 (Glutaredoxin), Solyc08g006250 (Copper transporter), Solyc07g049440 (GDSL esterase/lipase), Solyc02g062130 (Ferredoxin-NADP reductase) and Solyc08g007770 (PGR5-like protein 1A) were found. For *SlNRT2.2* gene, Solyc08g080340 (Lysophospholipid acyltransferase), Solyc04g082140 (Laccase-22), Solyc07g042170 (Jasmonate ZIM-domain protein 3), Solyc02g065470 (Pathogenesis-related protein) and Solyc09g092450 (Long-chain-fatty-acid CoA ligase) were identified. For *SlNRT2.3* gene, Solyc01g073820 (CHP-rich zinc finger protein-like), Solyc02g092250 (Cytochrome P450), Solyc01g009320 (U-box domain-containing protein 24), Solyc03g113040 (ATP-binding cassette (ABC) transporter 17) and Solyc01g014230 (U-box domain-containing protein 4) were obtained. For *SlNRT2.4* gene, Solyc05g008150 (Metallocarboxypeptidase inhibitor), Solyc10g018320 (Pectinesterase), Solyc09g008500 (Non-specific lipid-transfer protein), Solyc01g010730 (Serine carboxypeptidase) and Solyc10g054750 (Auxin-induced SAUR-like protein) were found. When the co-expression network is examined, the fact that the *SlNRT2* genes are associated with genes from many different molecular pathways (transport, photosynthesis, transcription factors, cell wall modification, fatty acid metabolism, amino acid catabolism, hormone response) may prove to be important in cellular metabolism.

**Fig. 6.**
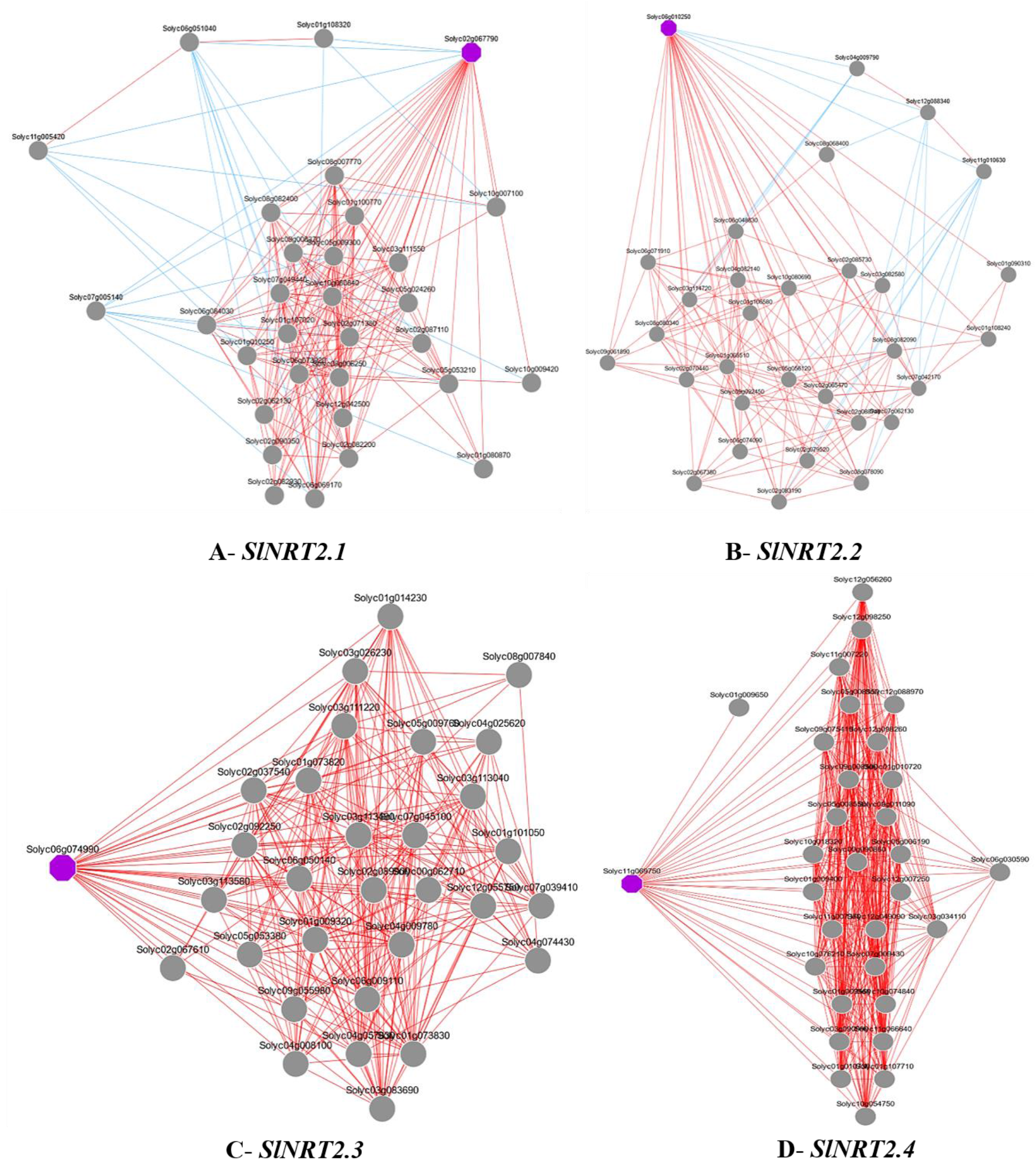
Co-expression network analyses of each *SlNRT2* genes indicated as violet color under abiotic stress condition using TomExpress platform based on RNA-seq data of vegetative and reproductive organs.

### Predicted 3D structure and phosphorylation sites

For seconder structure analyses, percentages of α-helix (47.39%, 47.99%, 45.25% and 44.76%), extended strand (16.30%, 15.06%, 18.25% and 15.81%), β-turn (5.43%, 7.83%, 6.27% and 6.10%) and random coil (30.87%, 29.12%, 30.23% and 33.33%) were identified for SlNRT2.1, SlNRT2.2, SlNRT2.3 and SlNRT2.4 proteins, respectively by using SOPMA online server. These variations in seconder structures may be related with functional diversities of *SlNRT2* genes in cell metabolism. When the predicted 3D structures of SlNRT2 proteins were examined, it was found that there were variances in both transmembrane helix numbers and channel numbers, which can be thought to increase the molecular diversity capacity of *SlNRT2* genes in cellular metabolism (Fig. 7).

**Fig. 7.**
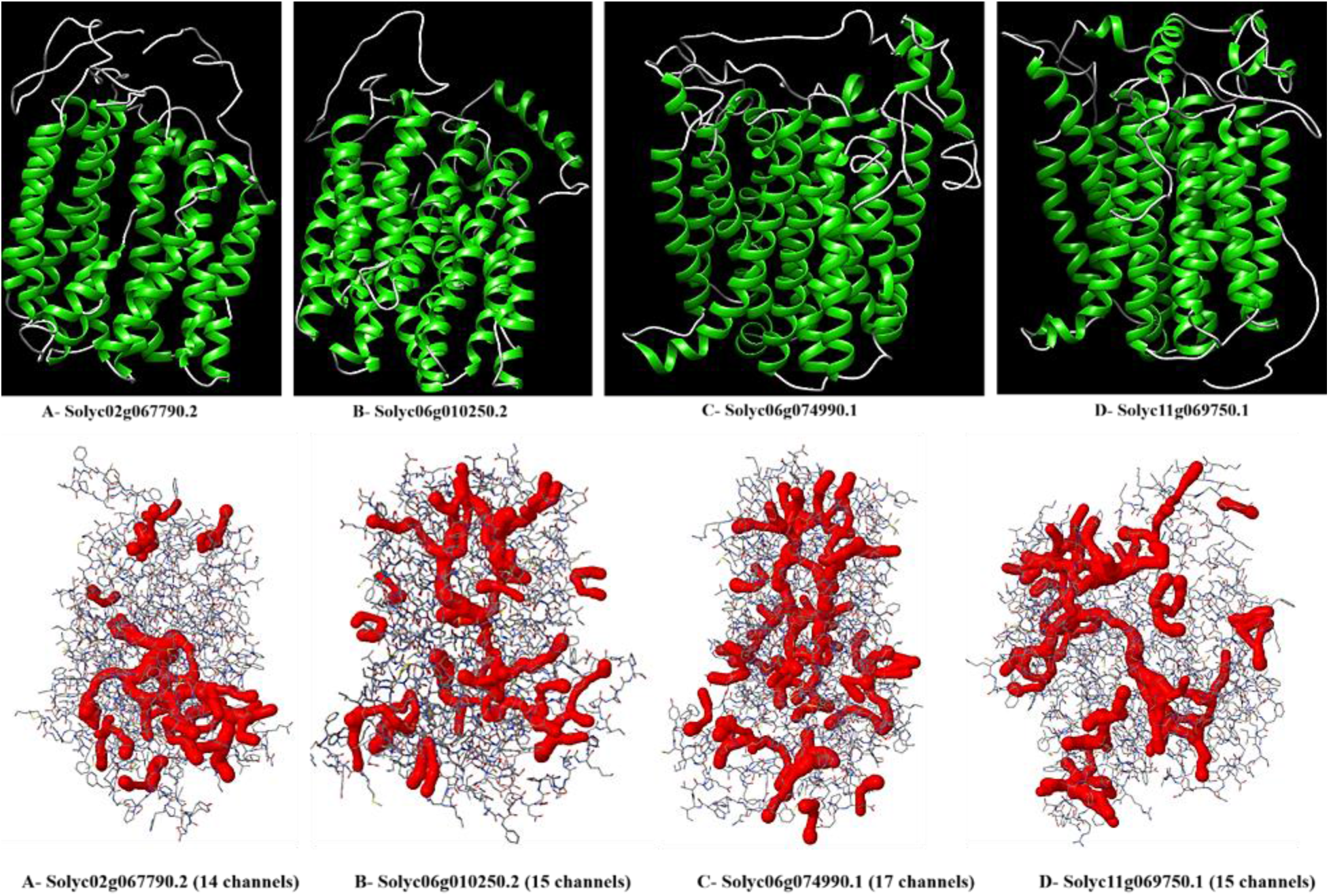
Predicted 3D structures and molecular cavities & their geometric properties of SlNRT2 proteins using Phyre2 and BetaCavityWeb server, respectively. The green and red structures show α-helices and channels respectively in SlNRT2 proteins.

The putative post-translational modifications of SlNRT2 proteins were identified using various online server, including serine, threonine or tyrosine phosphorylation, N-Glycosylation sites, sumoylation, lysine acetylation and palmitoylation (Table 2). Protein post-translational modifications (PTMs) are crucial and fastest way to response to changes in the environment at molecular level and PTMs also play important roles in regulation of many metabolic processes in plants (Arsova et al. 2018). Particularly, phosphorylation plays vital roles in programing of many physiological and metabolic pathways in plants, such as root growth (Zhang et al. 2016), carbon metabolism (Wu et al. 2014). In this context, PTMs detected in SlNRT2 proteins, especially the high number of predicted phosphorylation, may be related to the dynamic regulation of SlNRT2 proteins in cellular metabolism.

**Table 2.** The predicted post-translational modifications of SlNRT2 proteins

|  | NetPhos 3.1 | NetNGlyc 1.0 | GPS-SUMO 2.0 | GPS-PAIL 2.0 | CSS-Palm |
| --- | --- | --- | --- | --- | --- |
| Solyc02g067790.2 | 48 | 3 | 2 | 3 | 2 |
| Solyc06g010250.2 | 51 | 2 | 3 | 4 | 2 |
| Solyc06g074990.1 | 50 | 3 | 2 | 4 | 2 |
| Solyc11g069750.1 | 53 | 3 | 2 | 6 | 2 |

NetPhos 3.1: Serine, threonine or tyrosine phosphorylation, NetNGlyc 1.0: N-Glycosylation sites, GPS-SUMO: Sumoylation, GPS-PAIL: Lysine acetylation, CSS-Palm: Palmitoylation

### Phenotype, H_2_O_2_, MDA and Proline analyses

To confirm the efficacy of the stress applied, the plants were both phenotypically and physiologically examined. In the phenotypical examination, it was observed that the plants wilted under drought stress as a result of water deficit while the development of salt-stress applied plants was retarded (Fig. 8). The salt stress was also resulted in a visible thickening in the young leaves as a result of excess chlorophyll accumulation (Fig. 8).

**Fig 8.**
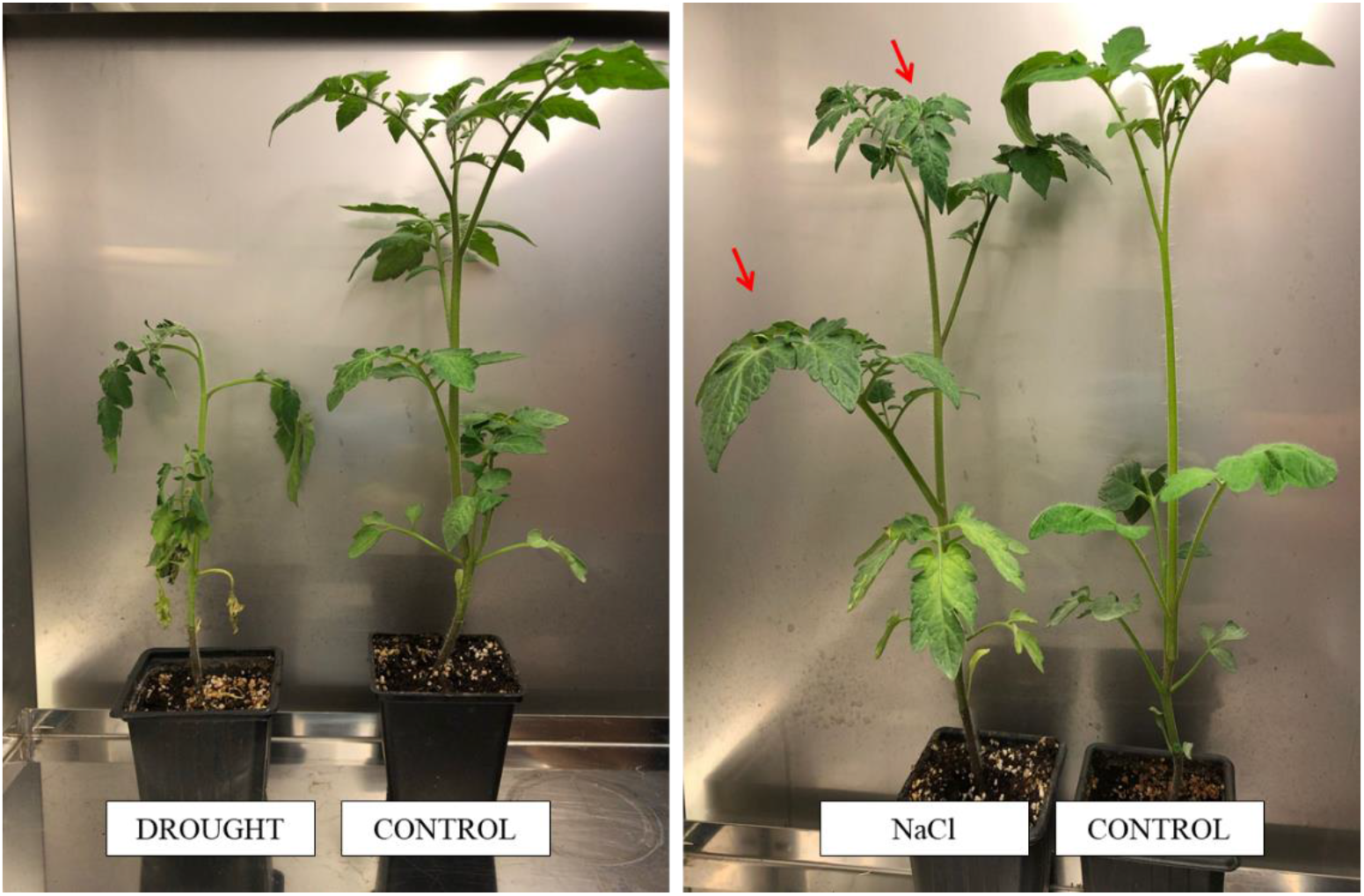
Effects of drought and salt stresses in tomato plants. The red arrows indicate excess chlorophyll accumulation and thickness in the leaves.

H_2_O_2_ is a byproduct of oxidative plant aerobic metabolism; therefore, it has been commonly used as an oxidative stress indicator (Saxena et al., 2016; García-Caparrós et al., 2020). Increase in H_2_O_2_ contents has been reported in drought- and salt-stressed tomato plants (Viveros et al., 2013; Jangid and Dwivedi, 2017). Additionally, proline content in the plants under the environmental stresses including drought and salinity stresses increase (Claussen 2005, Li et al., 2010). Generated as a result of lipid membrane peroxidation by ROS, MDA formation reflects stress-induced damage at cellular level (Marnett, 1999). Therefore, H_2_O_2_, proline and MDA contents of the plants are good environmental stress indicators to evaluate plant stress levels physiologically.

In the present study, H_2_O_2_ content was changed 2-fold in the drought-imposed plants leaves while no change was determined in their roots compared to the control (Fig. 9A). The NaCl stress exposure led to limited H_2_O_2_ accumulation in both roots and leaves, which were still important statistically.

**Fig 9.**
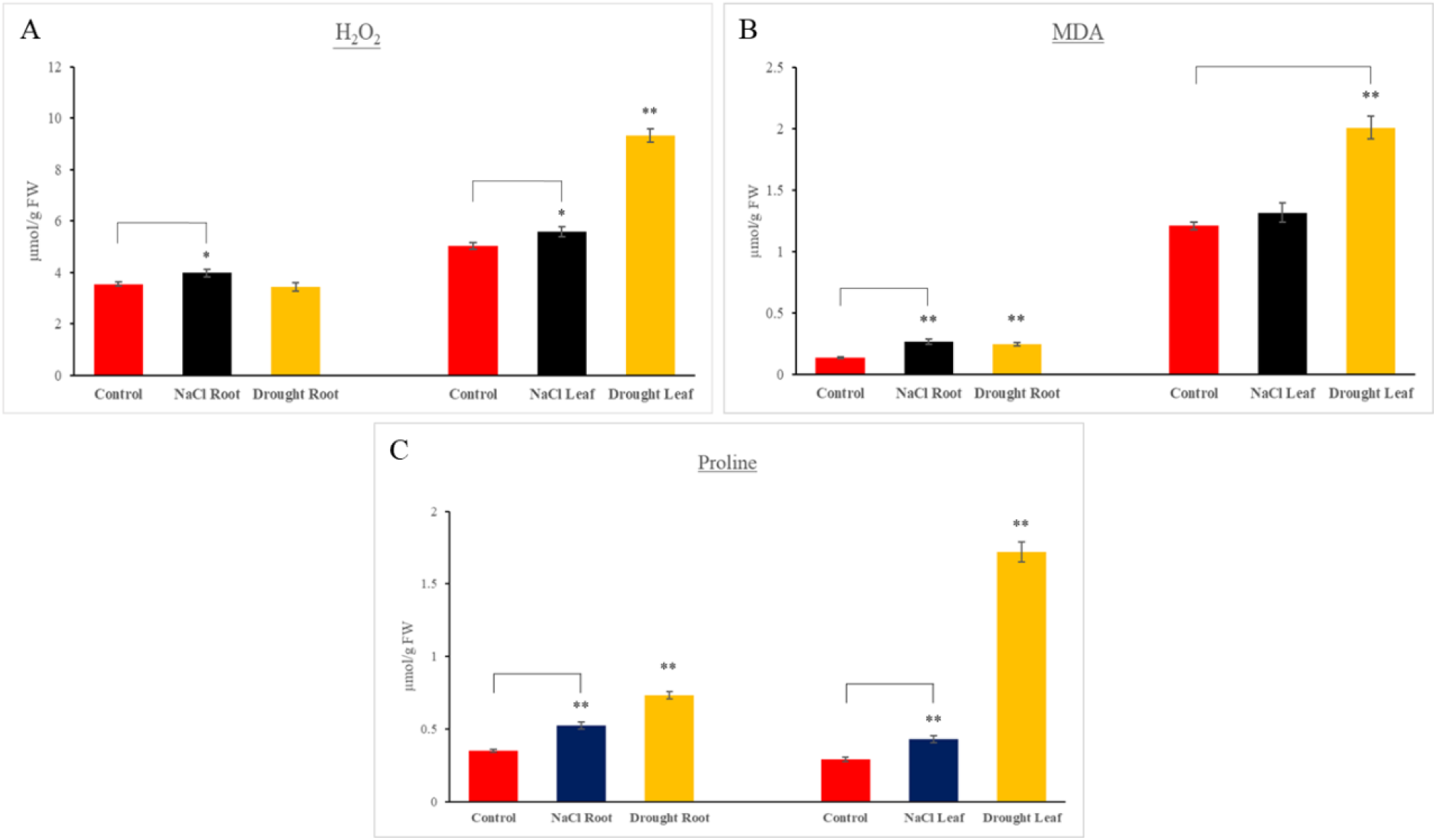
Effects of salt (NaCl) and drought stresses on H_2_O_2_ (μmol g^−1^ FW) (A), MDA (nmol g^−1^ FW) (B) and proline (μmol g^-1^ FW) (C) contents of tomato plants. Error bars depict standard errors of the mean (sdom; n = 3). Asterisks represent statistical significance (*: P > 0.05 and **: P > 0.01).

The MDA accumulation showed that drought caused a severe damage in both roots and leaves while NaCl stress only affected the roots (Fig. 9B). These results reveal that drought stress has a potent role on oxidative damage and lipid peroxidation in tomato cells. On the other hand, NaCl stress does not resulted in lipid peroxidation in tomato leaves.

A good stress indicator, the proline accumulation revealed that both drought and salinity applied led to stress in the plants non-organ-specific manner (Fig. 9C).

### *SlNRT2* gene expression analyses under salinity and drought stresses

## Author contribution statement

EF and MAA conceived the study, DC and EF conducted the experiments, and MAA and EF wrote the manuscript. The authors read, edited and approved the manuscript.

